# The longitudinal stability of fMRI activation during reward processing in adolescents and young adults

**DOI:** 10.1101/2020.08.06.236596

**Authors:** David AA Baranger, Morgan Lindenmuth, Melissa Nance, Amanda E. Guyer, Kate Keenan, Alison E Hipwell, Daniel S Shaw, Erika E Forbes

## Abstract

**Background:** The use of functional neuroimaging has been an extremely fruitful avenue for investigating the neural basis of human reward function. This approach has included identification of potential neurobiological mechanisms of psychiatric disease and examination of environmental, experiential, and biological factors that may contribute to disease risk via effects on the reward system. However, a central and largely unexamined assumption of much of this research is that neural reward function is an individual difference characteristic that is relatively stable over time.

**Methods:** In two independent samples of adolescents and young adults studied longitudinally (*Ns* = 145 & 153, 100% female & 100% male, ages 15-21 & 20-22, 2-4 scans & 2 scans respectively), we tested within-person stability of reward-task BOLD activation, with a median of 1 and 2 years between scans. We examined multiple commonly used contrasts of active states and baseline in both the anticipation and feedback phases of a card-guessing reward task. We examined the effects of cortical parcellation resolution, contrast, network (reward regions and resting-state networks), region-size, and activation strength and variability on the stability of reward-related activation.

**Results:** Overall, stability (ICC; intra-class correlation) across 1-2 years was modest. In both samples, contrasts of an active state relative to a baseline were more stable (e.g., Win>Baseline; mean ICC = 0.13 – 0.33) than contrasts of two active states (e.g., Win>Loss; mean ICC = 0.048 – 0.05). Additionally, activation in reward regions was less stable than in many non-task networks (e.g., dorsal attention), and activation in regions with greater between-subject variability showed higher stability in both samples.

**Conclusions:** These results show that functional neuroimaging activation to reward has modest stability over 1-2 years. Notably, results suggest that contrasts intended to map cognitive function and show robust group-level effects (i.e. Win > Loss) may be less effective in studies of individual differences and disease risk. The robustness of group-level activation should be weighed against other factors when selecting regions of interest in individual difference fMRI studies.

## Introduction

The translational relevance of neural reward processing research is evident from a large and growing literature revealing group differences in regional functional magnetic resonance imaging (fMRI) activation during reward processing across a range of neuropsychiatric conditions, including anxiety, ADHD, depression, addiction, schizophrenia, and autism (Chase et al., 2018; Clements et al., 2018; Forbes et al., 2006; Guyer et al., 2012; Luijten et al., 2017; Ng et al., 2019; Scheres et al., 2007). Building on this work, neural response to reward has been considered a potential biomarker (reflecting the presence of a disorder) or endophenotype (reflecting genetic risk) for these same conditions (Caseras et al., 2013; Dichter, 2012; Grimm et al., 2014; Hasler et al., 2004; Moeller and Paulus, 2018; Pizzagalli, 2014; Rubia, 2018; Sutherland and Stein, 2018). A major implication of this work is that reward-related neural activation may reflect causal neurobiological processes that underlie the emergence of neuropsychiatric disorders. Accordingly, a host of individual difference studies have sought to uncover potential mechanisms that may influence disease risk via their effects on reward processing, such as genetic risk and stress exposure (e.g. Banihashemi et al., 2014; Carey et al., 2017; Casement et al., 2015; Corral-Fríasa et al., 2015; Hanson et al., 2015b, 2015a; Jia et al., 2016; Kumar et al., 2015; Luking et al., 2016; Novick et al., 2018; Romens et al., 2015; Ruggeri et al., 2015). By examining possible causal mechanisms underlying mental illness, findings from this line of research have the potential to have a large impact on the development of novel treatment and prevention measures.

A central assumption of much of this work relating neural response to reward to the presence of, or risk for, a psychiatric disorder is that patterns of neural response to reward, as measured by fMRI, is a stable trait. However, only a few studies have examined the stability of neural response to reward stimuli over time. Two prior studies have used different versions of the monetary incentive delay (MID) task, in which participants are told whether the potential outcome of a trial is win, loss (Keren et al. did not include loss trials), or neutral, and then instructed to respond quickly to a cue (Keren et al., 2018; Wu et al., 2014). Wu et al. and Keren et al. both examined the longitudinal stability (e.g. over 2+ years) of activation within *a priori* ROIs using versions of the MID, in adults and adolescents (ns=14 & 16), respectively. Wu et al. tested the stability of anticipatory activation, and observed that the nucleus accumbens and insula were stable only during gain and loss anticipation, respectively (ICCs 0.5-0.7). In contrast, Keren et al. found stable activation only during reward outcome in the accumbens (peak ICC = 0.23) and a stable reward prediction error (RPE) signal in the insula (peak ICC = 0.16). Using a gambling reward Braams et al. examined the nucleus accumbens in the Win>Loss contrast during reward outcome (*N*=238, ages 8-26), and reported a similar stability estimate to Keren et al. (ICCs=0.219-0.327) (Braams et al., 2015). This work provides initial evidence that the nucleus accumbens has moderate stability during reward processing in both adolescents and adults. However, the impact of analytic decisions, such as choice of task phase (i.e. anticipation vs. feedback) and contrast (i.e. Win > Neutral vs. Win > Loss) remains unknown, this prior work may suffer from small sample sizes, and the stability of regions outside of the nucleus accumbens—even of regions known to play key roles in reward processing that have also been implicated in the etiology of neuropsychiatric disorders (e.g. the orbitofrontal cortex (Ng et al., 2019))— has been largely unexplored. As a result, there is limited evidence that these findings generalize.

Beyond this initial work examining the stability of reward activation, little is known about the factors associated with the reward activation stability. Candidate factors include region of interest (ROI) and region size, activation strength, activation variability, and network membership (e.g. reward, salience, or control). Although prior work has largely focused on regions considered to be part of the canonical reward network, reward tasks invariably elicit activation in other networks - such as in frontoparietal, cingulo-opercular, motor, and visual regions. Furthermore, activation of these regions may be less specific to reward task cues, but they still play important roles in reward function (Haber and Knutson, 2010; Schultz, 2000), and could contribute to differences in reward behavior between diagnostic groups. It is also clear that group-level contrast maps for reward tasks are highly reproducible (e.g., studies report activation of a consistent set of canonical reward regions) (Bartra et al., 2013; Jia et al., 2016; Kampa et al., 2020), suggesting that reward and non-reward regions may differ in their stability. Additionally, evidence suggests that the homogeneity of fMRI activation in a reward task varies as a function of ROI size (Schaefer et al., 2017); given that there are approximately 400 cortical areas (Van Essen et al., 2012), ROI size may influence stability of activation over time. As a whole, more information is needed about the stability of reward activation with regard to these various factors in order to advance longitudinal and clinical neuroscience research.

It is of critical importance to understand the influence of developmental timing on reward stability. Structural and functional brain maturation, including reward function, co-occurs with emergence or worsening of mental health problems during adolescence and young adulthood (Caspi et al., 2020; Foulkes and Blakemore, 2018; Galvan, 2010; Kessler et al., 2005; Kessler and Wang, 2008). Although capturing stability during this period of development is likely to be challenging given individual differences in the pace of maturation, such individual differences would be highly relevant to studies of disease risk, progression, and preventive intervention.

The present study aims to test the stability of individual differences in reward function measured during a widely-used card guessing task with monetary reward in two independent, adolescent and young-adult samples (*N*s=145 and 153 respectively). The current research represents the largest study of whole-brain stability in a reward fMRI task to date. We compared examined the stability of regional activation across the most commonly used contrasts from this task across the whole brain, and as a function of network identity, ROI size, and activation magnitude and variability. We focused on these aspects in relation to assessing stability of neural response to reward with the goal of informing the optimal analysis of fMRI reward tasks to be used in future individual difference studies. We hypothesized that the stability reward activation would be higher in young adult than adolescent samples, estimates of activation in canonical reward regions would be more stable over time than in regions within other networks, and that ROI size would be correlated with stability.

## Methods

Data were drawn from two independent longitudinal neuroimaging studies, the Pitt Mother & Child Project (PMCP) and the Pittsburgh Girls Study – Emotions Substudy (PGS-E), both administered the same reward processing task and were conducted with MRI scanning at the same location. In both studies, all procedures received Institutional Review Board approval at the University of Pittsburgh and all participants provided consent for their participation in the study.

### Pitt Mother & Child Project (PMCP)

The Pitt Mother & Child Project (PMCP) was a longitudinal study of 310 boys and their families residing in low income/resourced environments. Participants were recruited in 1991 and 1992 from Allegheny County Women, Infant and Children (WIC) Nutritional Supplement Clinics when boys were between 6 and 17 months old (Shaw et al., 2012, 2003). Boys and their mothers or primary caregivers were seen almost yearly from ages 1.5 – 22 years in the laboratory and/or home where they completed questionnaires, a psychiatric interview, and, at ages 20 and 22 years, fMRI scanning sessions. Study visits and interviews occurred as close to participants’ birthdays as was practically possible. Of the 310 participants, 139 had usable data available for both scans (see Table 1 for demographics). PMCP participants were excluded from the current study if (1) they could not be recruited for an MRI visit (*n*=75; i.e., refused or unable to participate), (2) they were ineligible for an MRI scan due to medical or physical reasons (*n*=49; i.e., concussion, recent drug use or metal in body), (3) they did not complete the MRI portion of the study (n=25), (4) they did not correctly perform the task (*n*=13; i.e., fell asleep, <80% accuracy, or did not understand instructions), or (5) the fMRI scan did not pass quality control benchmarks (*n*=10; e.g. excessive movement or poor coverage of the nucleus accumbens). The majority of participants were either White American (*n*=72, 51.8%) or Black/African American (*n*=55, 39.57%), with the remainder identifying as bi-racial (*n*=8, 5.76%), or Native Hawaiian, Native American, or Mexican American (*n*=4, 0.72%). These racial/ethnic demographics are consistent with city demographics at the time of recruitment.

**Table 1.** Sample demographics

|  |  |  |  |  |  |  |  |
| --- | --- | --- | --- | --- | --- | --- | --- |
| PMCP | N=139 (n=278 scans; 2 scans per participant) |  |  |  |  |  |  |
| Age by wave | Wave 1 | Wave 2 |  |  |  |  |  |
|  | 20.06 (0.21) | 22.04 (0.13) |  |  |  |  |  |
| Race | White American | African American | Mixed | Other |  |  |  |
|  | n=72 (51.8%) | n=55 (39.57%) | n=8 (5.76%) | n=4 (0.72%) |  |  |  |
| Time between scans (years) | Mean (SD) | Median (IQR) | Min | Max |  |  |  |
|  | 1.98 (0.23) | 2 (1.95-2.06) | 0.77 | 2.7 |  |  |  |
| PGS-E | N=145 (n=439 scans; 2-4 scans per participant) |  |  |  |  |  |  |
| Number of scans | 2 scans | 3 scans | 4 scans |  |  |  |  |
|  | n=45 (31.1%) | n=51 (35.17%) | n=49 (33.79%) |  |  |  |  |
| Age by wave | Wave 1 | Wave 2 | Wave 3 | Wave 4 |  |  |  |
|  | 16.85 (0.51) | 18.01 (0.5) | 19.18 (0.47) | 20.28 (0.46) |  |  |  |
| Age at scan | 15 yrs. old | 16 yrs. old | 17 yrs. old | 18 yrs. old | 19 yrs. old | 20 yrs. old | 21 yrs. old |
|  | n=7 (1.6%) | n=61 (13.9%) | n=101 (23%) | n=101 (23%) | n=101 (23%) | n=61 (13.9%) | n=7 (1.6%) |
| Race | White American | African American | Mixed |  |  |  |  |
|  | n=35 (24%) | n=101 (69.66%) | n=9 (4.83%) |  |  |  |  |
| Time between scans (years) | Mean (SD) | Median (IQR) | Min | Max |  |  |  |
|  | 1.27 (0.54) | 1.157 (0.99–1.33) | 0.44 | 4.18 |  |  |  |

Descriptive information of the sample demographics of the Pitt Mother & Child Project (PMCP) study and the Pittsburgh Girls Study – Emotions substudy (PGS-E). SD= standard deviation. IQR = inter quartile range (Q1 – Q3).

### Pittsburgh Girls Study – Emotions Sub-study (PGS-E)

Participants were girls and their mothers recruited from the longitudinal Pittsburgh Girls Study (PGS) (Keenan et al., 2010). The PGS sample was formed following an enumeration of households with girls between the ages of 5 and 8 in the city of Pittsburgh. Of the 2990 eligible families, 2450 (85%) were successfully re-contacted and agreed to participate in a prospective study. A subset of PGS participants was recruited to the PGS Emotions Sub-study (PGS-E) a study of precursors to depression (*N*=232 (Keenan et al., 2008)). From ages 15-21, participants were invited to complete four annual study visits and fMRI scanning sessions (see Table 1 for demographics). Of the potential *n*=928 scans (4 per participant), scans were unavailable or excluded from analyses because (1) participants withdrew or could not be scheduled (*n*=228 scans across *n*=121 participants), (2) participants were ineligible for an MRI scan due to medical or physical reasons (*n*=80 scans, *n*=55 participants), (3) participants refused or were unable to complete the MRI portion of the study (*n*=69,*n*=48 participants), (4) participants did not correctly perform the reward task or scans did not pass MRI quality control benchmarks (*n*=86, *n*=65 participants), or (5) because the aforementioned reasons resulted in only one usable MRI session (*n*=38 participants). The final sample consisted of 439 scans from *n*=145 participants, 49 participants had four scans, 51 had three scans, and 45 had two scans (Figure S1). The average time between scans was 1.27 years (SD=0.5), which ranged from 0.44 to 4.18 years. The majority of participants were Black/African American (*n*=101, 69.6%) or White American (*n*=35, 24%), with the remainder identifying as multi-racial (*n*=9, 4.8%).

### Monetary Reward and Loss fMRI Task

Participants in both samples completed the same version of an 8-minute slow event-related card-guessing task involving anticipation and receipt of monetary reward (Figure S1) (Forbes et al., 2010). In each trial (20 s), participants were shown a card with a question mark, told the possible value of the card was 1–9, and asked to guess whether the card’s value was lower or higher than 5 (4 s). Participants then learned the trial type (6 s) – possible-win or possible-loss (i.e., the anticipation phase). Participants were told the “correct” answer (500 ms) and then whether they had a positive (gain money), negative (lose money), or neutral (no-change) outcome (500 ms; i.e., the outcome phase). Each trial ended with a cross-hair that was presented during a 9 second inter-trial interval. Participants completed 24 trials, with a balanced number of trial types (i.e., 12 possible-win and 12 possible-loss). Trials were presented in a pseudorandom order with predetermined outcomes. Participants received $10 after completing the task.

### Neuroimaging data collection and preprocessing

#### Data collection

All participants were scanned using the same Siemens 3T Trio scanner. BOLD functional images were acquired with a gradient echo planar imaging (EPI) sequence and covered 39 axial slices (3.1 mm wide) beginning at the cerebral vertex and extending across the entire cerebrum and the majority of the cerebellum (TR/RE=2000/25ms, field of view =20cm, matrix=64×64). A reference EPI scan was acquired before fMRI data collection, which was visually inspected for artifacts (e.g., ghosting) and for adequate signal across the entire volume. A 160-slice, high-resolution, sagittally acquired T1-weighted anatomical image (MPRAGE) was collected for co-registration and normalization of functional images (TR/TE = 2300/2.98 ms, field of view = 20 cm, matrix = 256 × 240).

Preprocessing and within-person analysis: Preprocessing for both samples was completed using Statistical Parametric Mapping software (SPM8; http://www.fil.ion.ucl.ac.uk/spm). Structural images for each participant were auto-segmented, and functional images were realigned to correct for head motion, registered to the segmented structural data, spatially normalized into standard stereotaxic space (Montreal Neurological Institute template) using a 12-parameter affine model, and smoothed with a 6mm full-width at half-maximum Gaussian filter. Voxel-wise signal was ratio-normalized to the whole-brain global mean. The Artifact Detection Toolbox (ART; http://www.nitrc.org/projects/artifact_detect/) software was used to detect functional volumes with movement > 3 SD from the subject’s mean, > 0.5 mm scan-to-scan translation, or > 0.01 degrees of scan-to-scan rotation. Preprocessed data were inspected to ensure that all participants had fewer than 25% of volumes with excessive movement detected by ART, good scan quality, and ventral striatum coverage of at least 80%. Temporal censoring based on ART output was used to remove motion artifacts in first-level analyses.

The two research groups (PMCP and PGS-E) used slightly different time intervals within the course of the task to define the different phases of reward processing (e.g. anticipation and outcome). These differences have been consistent across the publications within each study (Casement et al., 2016, 2015, 2014; Hasler et al., 2017; Morgan et al., 2014; Romens et al., 2015). Thus, to maintain consistency with prior work and to compare the effects of differences in task phase definition within each study, no adjustments were made in the present analyses. In both studies, reward anticipation was defined as the 6 seconds when the symbol indicating trial-type was displayed, however reward outcome was defined differently: as the 1 second of outcome and the first 6 seconds of the inter-trial interval in the PMCP, and as the 1 second of outcome, and the first second of fixation in the PGS-E. The reward anticipation period extended 2 seconds beyond the potential-win arrow to account for the delay in hemodynamic response relative to neural activity and to capture as much of the reward anticipation response as possible while avoiding substantial overlap with BOLD response to reward outcome events. However, as the PGS-E definition of reward anticipation included the first 2 seconds of outcome (i.e., prior to the onset of the hemodynamic response to the outcome cue), the outcome phase was not included in analyses. For both studies, baseline was defined as the last 3 seconds of the inter-trial interval (during which a fixation was presented). First-level general linear models (GLMs) were used to calculate images for all contrasts, including: (1) win anticipation > loss anticipation, (2) win anticipation > baseline, (3) loss anticipation > baseline, (4) win outcome > loss outcome, (5) win outcome > baseline, (6) loss outcome > baseline, (7) win outcome > neutral outcome (e.g. no-win and no-loss), and (8) loss outcome > neutral outcome. Anticipation phase contrasts were calculated for both samples, while outcome-phase contrasts were calculated only for the PMCP.

Cortical regions of interest (ROIs) were defined using the Schaefer atlas (Schaefer et al., 2017), a recent cortical parcellation derived from resting-state fMRI data. One strength of this atlas, for the present analyses, is that parcellations at different resolutions (e.g., 100 – 1,000 regions) were identified, using identical methodology and data. Thus, the use of this atlas permits examination of the influence of ROI size, while holding potential confounds constant, including parcellation method (automatic vs. manual) and source data (histological vs. DTI vs. resting-state fMRI). To allow the broadest possible definition of regional response, all of the Schaefer atlas parcellations – 100, 200, 300, 400, 500, 600, 700, 800, and 1,000 ROIs – were used. Subcortical ROIs were identified using the Harvard-Oxford subcortical atlas (Frazier et al., 2005) using a threshold of 25% probability, and voxels present across multiple ROIs were discarded. All cortical and subcortical ROIs were included in analyses. Subject-level average percent signal-change was extracted from each ROI from every contrast in both samples, using the Marsbar toolbox for SPM (Brett et al., 2002).

### Statistical analyses

#### Stability

The longitudinal test-retest reliability, or stability (Becht and Mills, 2020), of the activation of each region in each contrast was estimated using the R-software package ‘RptR’ (Stoffel et al., 2017), in which a mixed-effect linear model was fit to all observations from all participants, with a random intercept for each participant. Participant race was added as a fixed effect, as were age and age^2^, as reward-related fMRI activation is known to vary as a function of age (Braams et al., 2015; Lamm et al., 2014). Regional activation was winsorized to 3 standard deviations prior to reliability estimation, to reduce the influence of outliers. Longitudinal reliability (stability) was then estimated as the ratio of the variance of within-person means over the sum of the group-level and residual variance. That is, reliability in the context of a longitudinal study is an estimate of the stability of individual differences over time (Revelle and Condon, 2018). As it is a ratio, this measure of stability (ICC_(3,1)_) is bounded by [0, 1]. Higher values, closer to 1, indicate that the measurement is more consistent over time. Models estimated the stability of each ROI in each sample, across all ROIs in every Schaefer parcellation (4,600 ROIs total across 9 parcellations) and the Harvard-Oxford atlas (14 ROIs), for all contrasts (8 contrasts in the PMCP, 3 in the PGS-E). *Post-hoc* analyses additionally examined the stability of *ranked* activation in the PMCP and PGS-E (Supplemental methods), as it is possible that there are group-level changes in mean activation beyond what is captured by participant age, which could bias stability estimates. The stability of ranked activation was strongly correlated with primary stability estimates across all parcellations, contrasts, and regions (PMCP: *r*=0.965, *p*<2.2×10^−16^; PGS: *r*=0.60, *p*<2.2×10^−16^). Results of these analyses are reported in the Supplement, as they do not meaningfully differ from the results and conclusions of the primary analyses.

#### Analyses

Linear mixed-effect models were used to test associations of parcellation, contrast, and ROI, entered as crossed random effects with random intercepts, with ROI stability. Stability estimates were winsorized to three standard deviations for these analyses, to reduce the influence of outliers. Variance explained by random effects was assessed using repeatability (R; i.e., the ratio of variance explained over total variance) calculated using the Rptr package (Stoffel et al., 2017). Once it emerged that parcellations did not differ in stability, and that the Win > Baseline contrast during the Anticipation phase had the highest stability (see Results), subsequent results considered only this contrast, using the *n*=400 Schaefer parcellation, given estimates of approximately 400 cortical regions (Van Essen et al., 2012), with the addition of subcortical ROIs. Analyses then examined the correlation between ROI stability and network identity (entered as dummy-variables), ROI size, average activation, and the variation of activation in both the PMCP and PGS-E. Average activation and the variation in activation of each ROI were estimated as the intercept and standard-error of the intercept in a linear mixed effect model with no fixed-effects, and participant as a random intercept. All analyses were conducted in R (R Core Team, 2014).

### ROI network assignment

The Schaefer parcellations include an assignment of each ROI to one of the seven canonical non-overlapping resting state networks: visual, somatomotor, dorsal attention, ventral attention, limbic, frontoparietal, and default mode (Yeo et al., 2011). However, these networks do not include subcortical regions, and there is no one network specific to the cognitive demands of the reward task used in the present analyses. Thus, an eighth ‘reward’ network was defined using Neurosynth (Yarkoni et al., 2011). Neurosynth is a platform that generates meta-analyses, using reported loci from published fMRI studies, thus identifying the regions of the brain most likely to be reported in studies on a given topic. We used the Neurosynth meta-analysis of reward-associated keywords, including “reward”, “outcome”, “anticipation”, and “monetary” (e.g., topic 7 from the v5 50-topic solution (Poldrack et al., 2012), which identifies voxels (*p*<0.01 false-discovery rate corrected) more likely to be reported in reward studies (*n*=1,218) than non-reward studies (*n*=13,153) (Supplemental Figure 3), https://neurosynth.org/analyses/topics/v5-topics-50/7. ROIs’ network-assignment was changed from Schaefer-assigned networks to the reward network, based on the overlap between voxels in the Neurosynth meta-analysis and each ROI. A *t*-statistic was calculated based on the distribution of this overlap and regions that showed a significant overlap (*p*<0.05) were assigned to the reward network. The reward network consisted of 22 regions, including the bilateral nucleus accumbens, caudate, putamen, pallidum, amygdala, and thalamus. Cortical regions included ROIs in the bilateral ventromedial prefrontal cortex and orbitofrontal cortex. The hippocampus was the only subcortical region not assigned to the reward network – it was manually assigned to the limbic network, as it has long been recognized as a central component of that system (Morgane et al., 2005). In analyses where networks were compared, the limbic network was set as the reference network, as it was observed to have the lowest stability (see Results).

## Results

### Parcellation resolution does not impact stability

Mixed effect models, in which parcellation was treated as a random intercept, revealed that stability did not differ across parcellations in the PMCP (R = 0, *p* = 1), but did in PGS-E. This minimal effect of parcellation (R=0.004, p=0.001) was driven by slightly lower stability in the 100- and 200-region parcellations (Figure S4). The addition of a second random intercept for task contrasts improved model fit and explained a significant amount of the variance of ROI stability (PMCP: R = 0.64, *p* < 2.2×10^−16^; PGS-E: R = 0.32, *p* < 2.2×10^−16^). There was no evidence for an interaction between parcellation and contrasts, as the addition of an interaction term did not improve model fit (PMCP: χ^2^_(1)_= 0, *p* = 1; PGS-E: χ^2^_(1)_= 1.9, *p* = 0.166) (Figure S5). The addition of a random intercept for each ROI explained a significant amount of the total variation in stability (PMCP: R = 0.11 *p* < 2.2×10^−16^; PGS-E: R = 0.266, *p* < 2.2×10^−16^), indicating that the stability of individual ROIs was somewhat consistent across contrasts. The stability of every ROI across all contrasts in both samples is provided in the Supplemental Data File.

### Reward anticipation and contrasts relative to baseline are more stable than loss contrasts or contrasts between active conditions

Given the evidence that stability did not vary as a function of parcellation resolution, analyses were then restricted to the Schaefer et al. parcellation with 400 parcels, based on estimates that there are approximately 400 cortical areas (Van Essen et al., 2012); subcortical regions were included in all subsequent analyses. In the PMCP, stability of average whole-brain activation was highest during reward anticipation in the Win > Baseline contrast (Figure 1A; 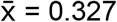, *SD* = 0.091). Win contrasts were associated with higher activation stability than Loss contrasts (β = 0.045, *SE* = 0.003, *t* = 15.71, *p* < 2.2×10^−16^), and contrasts relative to Baseline were associated with higher stability (β = 0.203, *SE* = 0.003, *t* = 70.77, *p* < 2.2×10^−16^) (Figure 1A, Table S1). The feedback phase was associated with lower activation stability than the anticipation phase (β = −0.015, *SE* = 0.003, *t* = −5.29, *p* = 1.31×10^−7^). In the PGS-E, which only examined the anticipation phase, activation estimates from the Win and Loss contrasts, each relative to Baseline, were similarly more stable than those from the contrast between these two active conditions (β = 0.064, SE = 0.003, *t* = 19.54, *p* < 2.2×10^−16^) (Figure 1B, Table S1). However, during anticipation the Win > Baseline contrast was slightly less stable, on average, than Loss > Baseline (β = −0.008, *SE* = 0.003, *t* = −2.65, *p* = 0.008; 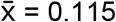, *SD* = 0.07).

**Figure 1:**
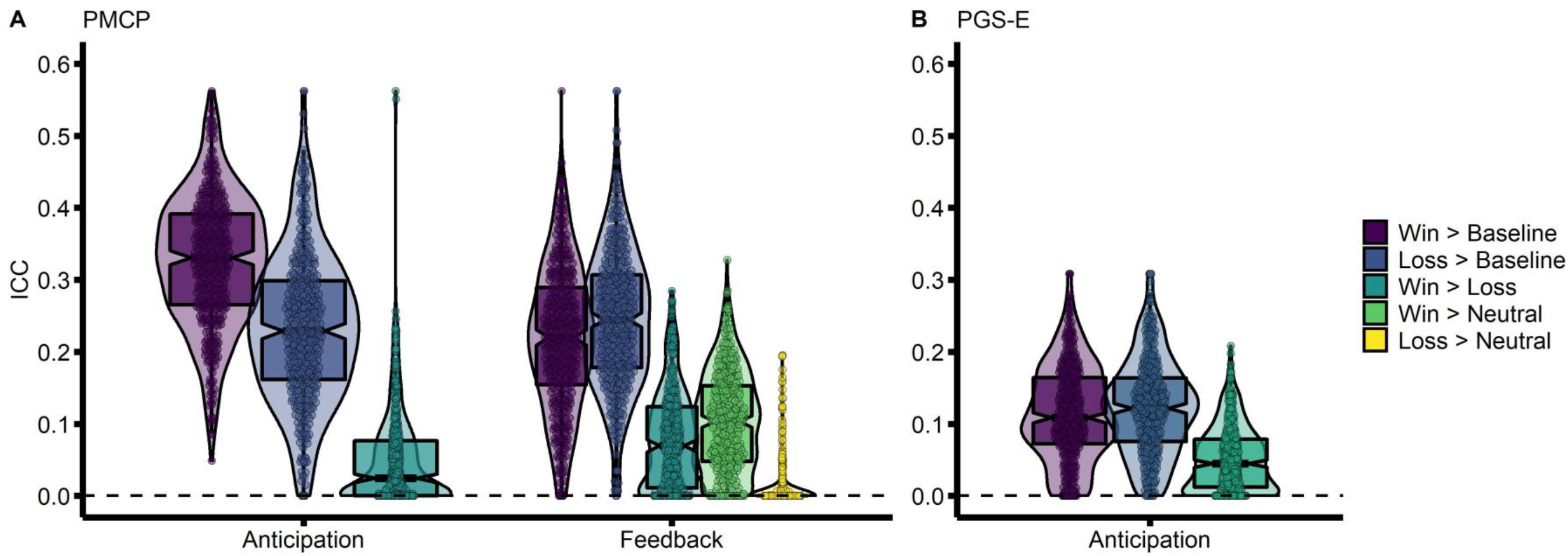
Win > Baseline contrast has a higher stability than Win > Loss contrast. Stability of activation across the whole brain, using the Schaefer-400 cortical parcellation. All five contrasts for examined task phases (anticipation and feedback) are shown. (A) The Pitt Mother & Child Project (PMCP; N=139) sample. (B) The Pittsburgh Girls Study – Emotions substudy (PGS-E; N=145) sample. Associated statistics are reported in the main text, as well as in Supplemental Table 1.

### Stability is lower in the limbic and reward networks than other cortical networks

Differences between the average stability of networks during the anticipation of rewards (Anticipation > Baseline) was examined next. In the PMCP, results from an ANOVA showed a significant effect of network (*F*_7, 406_ = 15.5, *p* < 2.2×10^−16^, adjusted R^2^ = 0.197). Relative to the limbic network, the reward network did not significantly differ, whereas for all others (i.e., frontoparietal, dorsal attention, salience, default mode, visual, and somatomotor) activation estimates were more stable (Figure 2A, Table S2 and S3). In the PGS-E, there was similarly a significant effect of network (*F*_7, 403_ = 11.47, *p* = 2.67×10^−13^, adjusted R^2^ = 0.152). The limbic network was also the least stable in the PGS-E (Figure 2B, Table S2), and all networks were more stable than it, except for the reward network, which was not significantly different (Table S3).

**Figure 2:**
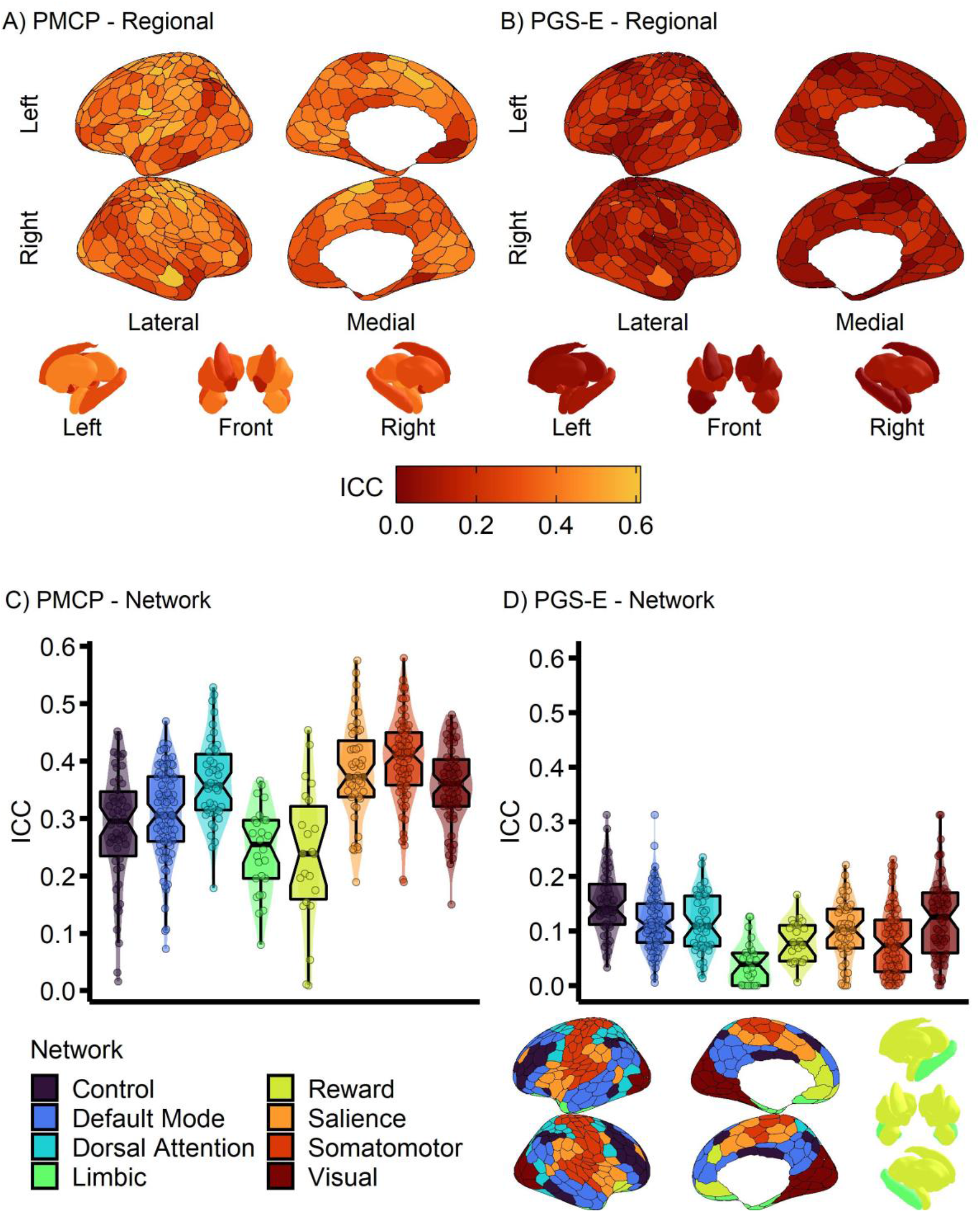
Stability is lower in the limbic and reward networks. Stability of activation during the anticipation of monetary gains (Win>Baseline contrast), using the Schaefer-400 cortical parcellation and Harvard-Oxford subcortical atlas, across all networks examined. (A&B) ROI reliability visualized on a surface projection. (A&C) the Pitt Mother & Child Project (PMCP; N=139) study, and (B&D) the Pittsburgh Girls Study– Emotions substudy (PGS-E; N=145). (C&D) Associated statistics are reported in Supplemental Tables 2 and 3.

### Regions with higher between-subject variability are more stable

Average between-subject activation and activation variability – the intercept and SE of the intercept from mixed effect models predicting the activation of each region – as well as region-size, were then added to the linear model predicting stability. In the PMCP, these terms significantly improved model fit (*F*_3, 403_ = 43.38, *p* < 2.2×10^−16^, change in adjusted R^2^ = 0.19). Average activation was associated with lower stability (β = −0.09, *SE* = 0.0168, *t* = −5.55, *p* = 5.16×10^−8^), whereas activation variability was associated with greater stability (β = 4.78, *SE* = 0.517, *t* = 9.26, *p* < 2.2×10^−16^) (Figure 3). ROI size (# of voxels) was nominally associated with stability in the full model (β = 5.6×10^−5^, *SE* = 2.5×10^−5^, *t* = 2.32, *p* = 0.026). In the PGS-E, these terms similarly significantly improved model fit (*F*_3,400_ = 53.89, *p* < 2.2×10^−16^, delta adjusted R^2^ = 0.24). As in the PMCP, activation variation in the PGS-E was associated with greater stability (β = 5.745, *SE* = 0.455, *t* = 12.617, *p* < 2.2×10^−16^). Average activation was not associated with stability (β = 0.002, *SE* = 0.001, *t* = 1.37, *p* = 0.17), and ROI size was similarly nominally associated with stability (β = 4.2×10^−5^, *SE* = 1.8×10^−5^, *t* = 2.235, *p* = 0.026) (Figure 3). The replicable and strong association of activation variation with stability indicates that regions with greater between-person variation also tend to be the most stable over time.

**Figure 3:**
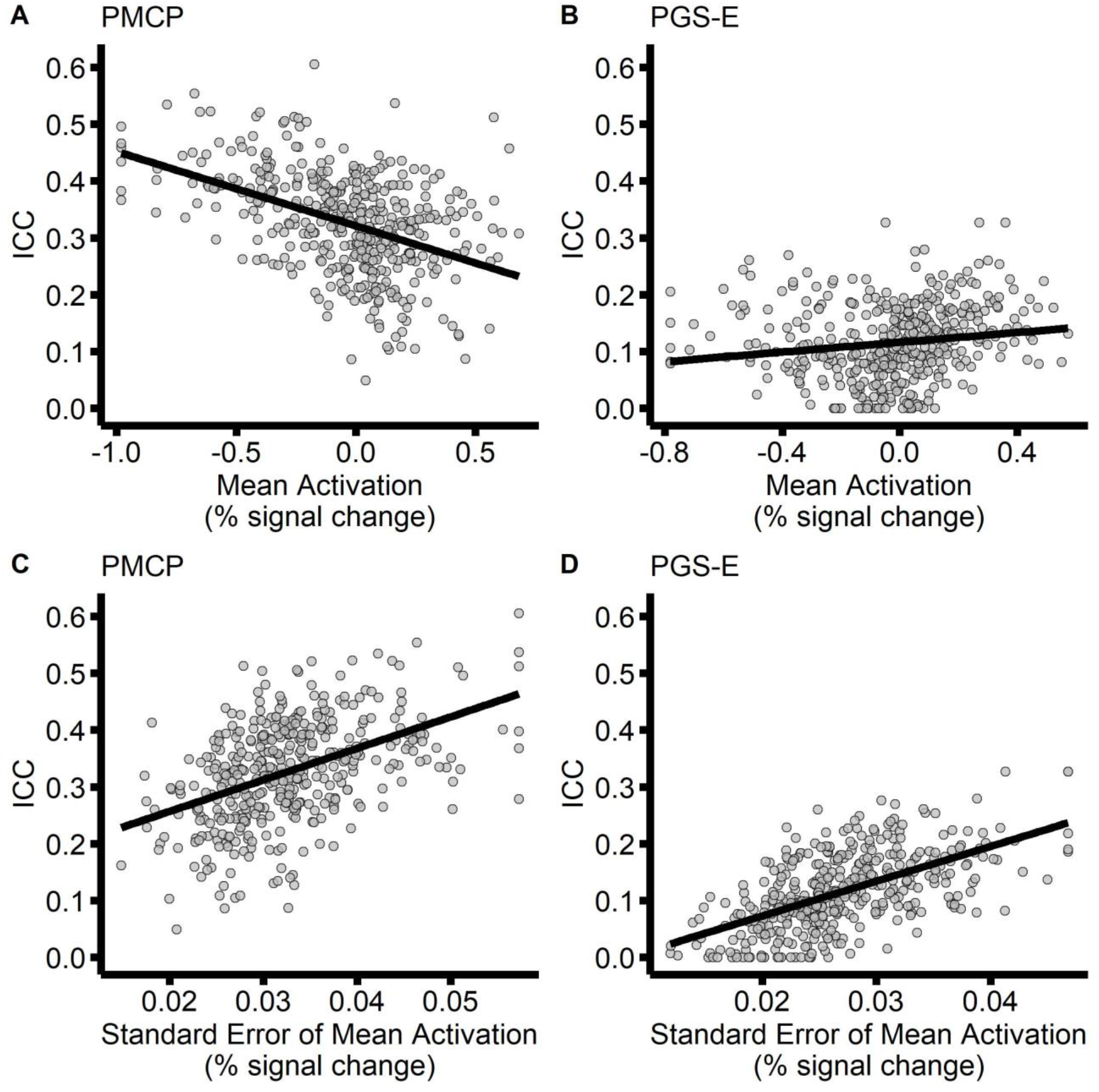
Regions with greater between-subject variation are more stable. Correlations between the stability of activation during the anticipation of monetary gains (Win>Baseline contrast), and region mean activation (A&B) and variability of activation (C&D). (A&B) the Pitt Mother & Child Project (PMCP; N=139) study, and (C&D) the Pittsburgh Girls Study– Emotions substudy (PGS-E; N=145).

To further assess the agreement in results between the two samples, the linear models correlating stability of activation estimates during the anticipation phase for the Win > Baseline contrast were used to predict stability across samples. Models included network identity, ROI size, and the mean and standard deviation of activation. Stability in the PMCP was predicted using results from the PGS-E, and conversely stability in the PGS-E was predicted using results from the PMCP. The predicted and true stabilities were significantly and moderately correlated in both samples (PMCP: *r*=0.38, r^2^=0.14, *p*=1.9×10^−15^; PGS-E: *r*=0.35, r^2^=0.12, *p*=1.7×10^−13^). This further indicates agreement in factors that influence stability, despite differences between the samples.

## Discussion

Our goal in the present study was to contribute data on the stability of reward function as measured by fMRI. This is one of the first studies to examine stability in relatively large samples of males and females, over a 1-2 year interval, and during the transition from adolescence to early adulthood. Our findings replicate results from the few existing studies; some reward contrasts, particularly contrasts of an active state against a baseline, are relatively stable over 1-2 years during this developmental period. In particular, while the magnitude of activation in many regions during the *anticipation* of reward was stable, few regions were stable when contrasting *win* relative to *loss*, a widely used contrast in studies of reward processing. Surprisingly, although we observed modest stability for the core reward-processing network, some of the highest stability estimates were found outside of that network. In addition, regions with the greatest between-person variability tended to show the highest level of stability.

### Contrasts of active conditions are less stable

One of the most striking observations from the present study is that contrasts of two active conditions (e.g. Win > Loss) were less stable than contrasts of a task condition to a fixation baseline. This was true in two independent samples, and in the PMCP the pattern was evident for both anticipation and outcome contrasts. Thus, we suggest that contrasts of two active task conditions, particularly of the reward task used here and similar tasks, may not be suitable as biomarkers for psychopathologies and may not reflect genetic or environmental risk for psychopathology, as they demonstrate poor stability. Thus, between-person differences in such contrasts may not be attributable to the presence of underlying traits.

The limited stability of Win > Loss activation in the present study adds to a growing body of work highlighting concerns over the reliability of fMRI task measures in general (Elliott et al., 2020; Fliessbach et al., 2010; Infantolino et al., 2018). The common practice of defining ‘activation’ as the difference in the average percent signal change between two different task states may contribute to the low reliability and stability (Luking et al., 2017). fMRI task activations computed this way are difference-scores; the reliability of a difference-score is partially a function of the correlation between the items being subtracted, and is lower for items that are more strongly correlated (Thomas and Zumbo, 2012). When two conditions, such as the anticipation of uncertain monetary *gain* or *los*s, are strongly correlated, the resulting difference score will have poor reliability.

Indeed, we note that studies of short-term reliability of reward activation yield stronger estimates when the contrast included a neutral comparison. Several studies have reported a moderate-to-good short-term (days or weeks) reliability of activation of the ventral striatum (ICC = 0.5 - 0.8) when examining the anticipation of monetary gain relative to a neutral condition where neither gain or loss was possible (Holiga et al., 2018; Plichta et al., 2012; Schlagenhauf et al., 2008). The neutral condition used in these studies differs from the neutral condition in the present study, a no-change outcome (for the PMCP sample only), which might be interpreted as a positive or negative outcome, depending on whether the trial involved not-losing or not-winning, respectively. In contrast, other studies have examined the short-term reliability of gain vs. loss contrasts, observing lower reliabilities in the striatum (ICC = −0.13 – 0.45) (Elliott et al., 2020; Fliessbach et al., 2010; Li et al., 2020). These results are congruent with our own results, suggesting that fMRI contrasts will be less stable if the cognitive processes underlying trial-types are too similar.

### Additional factors influencing stability

Beyond comparing contrasts, other factors that influence longitudinal stability were identified and found to be consistent across the samples used in this study, particularly the observation that greater between-subject variability of activation was associated with increased stability, and that ‘task specific’ ROIs tended not to be the most stable. The observation that more variable between-subject activation—in other words, regions that exhibited a greater range of individual differences—was predictive of greater longitudinal stability is not unique to this study. This pattern has been reported previously in different tasks (Fröhner et al., 2019) and modalities (resting-state (Pannunzi et al., 2017)), and has been discussed at length in the psychometric literature (e.g., the reverse, that effects with less between-subject variation are less reliable is sometimes referred to as the ‘reliability paradox’ (Hedge et al., 2018)). This observation in the present study may very well be a consequence of the computation of fMRI activation as a difference-score -- activation to different task conditions will be less correlated in regions with greater between-person variation, resulting in higher stability.

Interestingly, we observed that activation measured from task-specific ROIs tends to be less stable than that from non-task ROIs. In the present study, ‘task-specific’ ROIs were located primarily in the subcortex and ventral surface of the cortex, regions which are well-known to suffer from increased artifact susceptibility (Merboldt et al., 2000; Wiggins et al., 2009), which may reduce stability. Additionally, although the majority of studies that have examined the reliability or stability of reward task activation restricted their analyses to *a priori* reward-network ROIs, some looked beyond these structures. These studies have found that it is not uncommon for other regions of the brain to exhibit greater stability than the regions targeted by a task (i.e., regions in the default mode network have been frequently observed to be highly reliable) (Elliott et al., 2020; Fröhner et al., 2019; Holiga et al., 2018; Keren et al., 2018; Vetter et al., 2017). It is possible that regions performing computations not directly manipulated by the task may exhibit more stable inter-trial activation patterns, resulting in increased stability. These outcomes may include processes such as maintaining task goals and instructions (Dosenbach et al., 2006), or interpreting visual information. While the demands placed on these systems are relatively constant across the duration of a task, tasks are designed to elicit varied and pronounced activation in target regions – the reward system, in the case of the present study. Indeed, this feature is part of how these tasks produce robust and reproducible group-level activation maps. Yet at the same time, a varied pattern of activation may result in a less precise estimate of an individual region’s response in each participant, resulting in lower stability.

Several factors were not predictive of stability, including activation strength and region size. The finding that the overall strength of activation was not predictive of stability is consistent with our observation that regions within the reward network were not the most stable, and is consistent with prior results (Fröhner et al., 2019). As has been previously shown in other tasks, robust group-level between-condition differences do not imply that a measurement is stable or reliable (Infantolino et al., 2018). The present analyses used a set of cortical parcellations (Schaefer et al., 2017) that ranged in the number of regions, from 100 to 1,000. Doing so allowed us to explore the possible effects of ROI size and parcellation resolution, independent of potential confounding effects of parcellation method (e.g., DTI vs histology vs resting state fMRI). These effects were inconsistent across studies and in opposing directions. The parcellations with the largest ROIs (the 100 and 200 parcellations) were less stable in the PGS-E, but there was no effect on stability in the PMCP. In contrast, ROI size was very weakly associated with increased stability in the Schaefer-400 parcellation both samples. These results suggest that the choice of larger or smaller ROIs will not impact the sensitivity of individual-difference analyses, and that ROI choice should be guided by other considerations. For instance, smaller ROIs allow for the examination of more fine-grained effects on activation. However, this issue needs to be balanced with the consequent cost of a larger correction for multiple comparisons, which results in reduced power.

### Factors that may limit the maximally observable stability

Although we found that contrasts of neural activation to active condition relative to baseline were more stable than contrasts between different task conditions, observed stabilities were not in the range that is typically considered “optimal” (e.g., ICC > 0.7) (Cicchetti, 1994). However, this standard comes primarily from studies of inter-rater and short-term reliability. Studies of psychiatric diagnoses and psychopathology in adolescents and adults have reported a wide range of stabilities (Blázquez et al., 2019; Olino et al., 2018; Pettit et al., 2005; Shankman et al., 2017), suggesting that an ICC greater than 0.7 is not necessary for a measurement to reflect meaningful individual differences, but there is no currently accepted standard for longitudinal fMRI studies. Prior work has observed that fMRI activation can be within this range when certain contrasts are used (see the Discussion above), yet the same benchmarks may not be appropriate for stability across several years. Notably, prior meta-analyses have found that the reliability of both fMRI activation (Bennett and Miller, 2010) and resting-state correlations (Noble et al., 2019) decrease with longer intervals between scans. As an increasing number of longitudinal studies of brain function are conducted, it will be critical to report the stability of these measures, so that appropriate benchmarks can be determined (Herting et al., 2018).

Based on the large body of literature detailing the influence of development, aging, and life experience on brain function, it would be surprising if the longitudinal stability of neural activation estimates were as high as its short-term stability, especially during periods of known developmental change in the circuitry of interest (Braams et al., 2015; Galvan et al., 2006). Indeed, emerging work has suggested that the longitudinal stability of activation in feedback-learning and rule-learning fMRI tasks is highest in adult populations (Koolschijn et al., 2011; Peters et al., 2016). In the current study, the age of both samples provided the developmental window of late adolescence to early adulthood (15-22 years, inclusive) within which to measure brain function. This approach may have also contributed to the relatively lower observed stability. However, we note that the observed range of stabilities in the present analyses are comparable to longitudinal stabilities observed in adolescent samples performing other tasks, including cognitive control (Vetter et al., 2017), performance monitoring (Koolschijn et al., 2011), rule-learning (Peters et al., 2016), and emotional face monitoring (van den Bulk et al., 2013). Examining the influence of development on the stability of fMRI activation will be an important direction for future work.

We also should note that many additional factors likely influence the stability of neural activation as measured by fMRI tasks. Evidence suggests that activation during cognitive and affective tasks is sensitive to time-varying states, including mood, sleep, and recent stress (Nikolova and Hariri, 2012; Hasler et al., 2012; Baranger et al., 2016, 2017), which may place a ceiling on the maximum stability that is achievable with task-based fMRI. This concern is in addition to the wide array of technical aspects of fMRI data collection that will influence the signal-to-noise ratio, including task design, scanner manufacturer, ambient temperature, and head motion (Greve et al., 2011; Karch et al., 2019; Petersen and Dubis, 2012; Power et al., 2014).

### Comparisons between samples

One of the major strengths of the present report is the consistency of results across two samples that differed in several respects. Participants in the PMCP were all male, while participants in the PGS-E were all female. The ages of the two samples only partially overlapped – PGS-E participants (ages 15-21) were younger than PMCP participants (ages 20-22). However, there were some differences in the results from the two samples, particularly the lower overall stability in the PGS-E. There are several factors that may have driven this difference, including the age difference, differences in sample demographics, and differences in study methodology.

It is well documented that reward-related neural and behavioral processes undergo developmental changes into the 20s (Casey et al., 2008), and participants in both samples were likely undergoing development in reward and cognitive control circuitry, both of which mature during this period. Although the effects of developmental processes on stability are largely unknown, it is plausible that they reduce overall stability, as the effects of development are not uniform. Individuals’ trajectories of change occur at different times and rates, which would be expected to reduce the stability of measurements taken prior to, or during, the developmental period. The PMCP sample was entirely male, while the PGS-E was entirely female. Prior reports have observed greater activation during reward processing in males than females (Alarcón et al., 2017; Spreckelmeyer et al., 2009), which could be attributable to differential effects of sex hormones on reward sensitivity (Dreher et al., 2007; Harden et al., 2018). The samples also differed by socioeconomic status. All the participants in the PMCP came from families that were receiving economic benefits for low-income families (WIC) at the time of recruitment. The PGS over-sampled homes in low-resourced neighborhoods in the enumeration, resulting in approximately 40% of participants coming from families that received public assistance (Keenan et al., 2010). As a result, monetary rewards may not have had the same saliency across samples, which could produce differences in reward activation (Zink et al., 2004), resulting in stability differences.

The PMCP and PGS-E also took different approaches to scheduling participants. PMCP participants completed fMRI sessions as close to their 20^th^ and 22^nd^ birthdays as was practical, resulting in very little variation in the time between measurements (Supplemental Figure 1). The PGS-E took the more common approach of allowing flexibility in scheduling participants for their annual visit. Additionally, as the PGS-E collected fMRI data over a longer period, participants could miss one follow-up, but return for the subsequent visit, which was not possible in the PMCP, which included only two waves of fMRI data collection. These differences in study design resulted in much more variation in the time between measurements in the PGS-E, which may have contributed to the lower stability seen in the PGS-E sample. The additional waves of data collection in the PGS-E may have also contributed to the lower stability by virtue of reduced task novelty, though whether detectable habituation effects are present in fMRI measurements separated by years remains a largely unexplored topic (Telzer et al., 2018). Finally, the two studies differed in their definition of the anticipation phase of the task. In the PMCP, this phase ended once the feedback cue was presented, whereas in the PGS-E it was extended to include the feedback cue presentation, to account for the delay in the hemodynamic response to the anticipation cue. It is possible that activation during the extended anticipation phase may have been contaminated with activation reflecting subsequent processing.

### Future directions

The present study compared the stability of commonly used contrasts derived from an fMRI reward task, yet further work is needed to establish benchmarks for stability. Meeting these benchmarks may require the development of new tasks or paradigms, such as naturalistic imaging (Gruskin et al., 2019), or analyzing current tasks in different ways. For example, some reports suggest that correlations between trial-wise participant behavior (e.g., subjective value) and neural activation have increased reliability, relative to contrasts of activation in response to task conditions (Keren et al., 2018), but see also Fliessbach (Fliessbach et al., 2010). One intriguing recent report suggests that the low reliability of classic cognitive behavioral tasks can be resolved with hierarchical models that account for variability in trial-level behavioral responses, as opposed to simply averaging behavior across trials (Haines, 2019). A recent application of this approach to neuroimaging data suggests that it results in increased power (Chen et al., 2020). Related work has found that latent variable modeling can be used to improve the ability of fMRI analyses to detect individual differences (Cooper et al., 2019). Combining these approaches may prove to be a fruitful area of future research, particularly as they could be used to improve analyses of existing data sets.

## Conclusions

The findings from the present study suggest that in late adolescence through early adulthood, neural activation derived from a reward fMRI task has generally modest but acceptable stability over 1-2 years, with activation estimates in a contrast of two active conditions less stable than in a contrast of an active condition to a baseline. This difference likely accounts for some of the discrepancies in prior reports on the stability of fMRI activation. In addition, regions with greater between-subject variability and regions within non-task networks exhibited activation with greater stability, indicating that the robustness of group-level activation is not a sufficient sole criterion for selecting brain regions to use in individual difference studies. Further work is needed to establish benchmarks for the stability of fMRI tasks, and to develop tasks and analytic pipelines optimized for improving stability of neural activation measured with fMRI tasks. In sum, activation during reward fMRI tasks may be useful as a measure of stable individual differences for interrogating the etiology and course of psychopathology.

## Acknowledgements

The authors would like to thank the participants from both studies. This work was supported by grants from the National Institutes of Mental Health, including DA026222 (Shaw, Forbes), MH093605 (Guyer, Keenan, Forbes), and MH018951 (Baranger).

