## Supplemental Information for "The longitudinal stability of fMRI activation during reward processing in adolescents and young adults"

**Supplementary Information Table of Contents:**

Supplemental Methods: Page 3

Supplemental Results: Pages 4-6

Supplemental Tables 1 – 3: Pages 7-9

Supplemental Figures 1-4: Pages 10-14

References: Page 15

**Supplemental Methods**

Ranked Stability

To compute the long-term stability of ranked activation, activation estimates were ranked separately for each wave of the PMCP. For rankings to be meaningful in the PGS-E, a subset of scans were selected for analysis. Given the accelerated longitudinal design of the study, rankings of every scan would lose meaning as participants dropped-out and dropped-in at each subsequent wave. Scan-pairs whose time-between scan were within the interquartile-range of all scans (Table 1. Time-between scans = 0.99 – 1.33 years) were selected. When more than one scan-pair met this criterion for a participant, the pair whose interval was closest to group median (1.157 years) was selected. Scans from 100 participants (2-scans per participant) were included in the analysis. The earlier of the two scans for each participant was treated as the ‘first’ wave, and the later as the ‘second’ wave. Activation estimates within each wave were ranked separately. Estimation of stability was then conducted as described in the main Methods.

### Supplemental Results

Parcellation resolution does not impact long-term stability of ranked activation

Long-term stability did not differ across parcellations in either the PMCP (R = 0, $p$ = 1) or the PGS-E (R=0, p=1). As in primary analyses, the addition of a second random intercept for task contrasts improved model fit and explained a significant amount of the variance of ROI stability (PMCP: R = 0.65, $p < 2.2x10^{-16}$; PGS-E: R = 0.313, $p < 2.2x10^{-16}$). There was no evidence for an interaction between parcellation and contrasts, as the addition of an interaction term did not improve model fit (PMCP: $\chi^2_{(1)} = 0$, $p$ = 1; PGS-E: $\chi^2_{(1)} = 2.6$, $p$ = 0.11) (Figure S5). The addition of a random intercept for each ROI explained a significant amount of the total variation in stability (PMCP: R = 0.107 $p < 2.2x10^{-16}$; PGS-E: R = 0.267, $p < 2.2x10^{-16}$), indicating that the stability of individual ROIs was somewhat consistent across contrasts. The long-term stability of the ranked activation of every ROI across all contrasts in both samples is provided in the Supplemental Data File. We note that the effects reported here are nearly identical to those reported in primary analyses.

Reward anticipation and contrasts relative to baseline are more reliable than loss contrasts or contrasts between active conditions – ranked activation

In the PMCP, Win contrasts were associated with higher stability than Loss contrasts (β = 0.041, *SE* = 0.003, $t$ = 15.67, $p < 2.2x10^{-16}$), and contrasts relative to Baseline were associated with higher stability (β = 0.21, *SE* = 0.003, $t$ = 79.81, $p < 2.2x10^{-16}$). In contrast to the results of primary analyses, the Anticipation and Feedback phases did not differ (β = -0.002, *SE* = 0.003, $t$ = -0.94, $p$ =0.347). In the PGS-E, which only examined the anticipation phase, contrasts relative to Baseline were similarly more reliable than contrasts of two active conditions (β = 0.03, SE = 0.005, $t$ = 5.5, $p = 5.12x10^{-8}$). However, the Win Anticipation > Baseline contrast was less stable than Loss Anticipation > Baseline (β = -0.035, *SE* = 0.005, $t$ = -6.34, $p = 3.89x10^{-10}$). Comparing the stability of ranked activation to primary analyses within the Anticipation > Baseline contrast of the Schaefer-400 parcellation + subcortex found no difference in stability in the PGS-E ($t$=1.27, $p$=0.2), while in the PMCP the stability of ranked-activation was significantly lower ($t$=14.9, $p<2.2 \times 10^{-16}$; original mean = 0.327, ranked mean = 0.29).

Long-term stability of ranked activation by network

Differences between the average stability of networks during the anticipation of rewards (Anticipation > Baseline) were examined next. In the PMCP, results from an ANOVA showed a significant effect of network ($F_{7, 406}$ = 15.26, $p < 2.2 \times 10^{-16}$, adjusted $R^2$ = 0.195). Relative to the limbic network, the reward and default mode networks did not significantly differ, whereas all others (i.e., frontoparietal, dorsal attention, salience, visual, and somatomaotor) were more reliable. In the PGS-E, there was similarly a significant effect of network ($F_{7, 403}$ = 15.72, $p <$ $2.2 \times 10^{-16}$, adjusted $R^2$ = 0.2). In contrast to the PMCP, and results from primary analyses, the somatomaotor network was the least reliable. The reward network, as well as both the salience and dorsal attention networks did not differ from the limbic network.

Regions with higher between-subject variability are more reliable

Average between-subject activation and activation variability – the intercept and SE of the intercept from mixed effect models predicting the activation of each region – as well as region-size, were then added to the linear model predicting stability. In the PMCP, these terms significantly improved model fit ($F_{3, 403}$ = 21.32, $p < 2.2 \times 10^{-16}$, change in adjusted $R^2$ = 0.14). Average activation was associated with lower stability (β = -0.096, $SE$ = 0.0186, $t$ = -5.16, $p$ = $3.93 \times 10^{-7}$), while activation variability was associated with greater stability (β = 0.328, $SE$ = 0.045, $t$ = 7.2, $p$ = $2.8 \times 10^{-12}$). ROI size (# of voxels) was nominally associated with stability in the full model (β = $6 \times 10^{-5}$, $SE$ = $2.9 \times 10^{-5}$, $t$ = 2.13, $p$ = 0.03). In the PGS-E, these terms similarly significantly improved model fit ($F_{3,400}$ = 26.64 $p$ = 9.9x10$^{-16}$, delta adjusted $R^2$ = 0.13). As in the PMCP, activation variation in the PGS-E was associated with greater stability (β = 0.295, *SE* = 0.045, $t$ = 6.485, $p$ = 2.63x10$^{-10}$). Average activation was associated with increased stability (β = 0.0122, *SE* = 0.002, $t$ = 4.98, $p$ = 9.69x10$^{-7}$), and ROI size was weakly associated with stability (β = 1.44x10$^{-4}$, *SE* = 3x10$^{-5}$, $t$ = 4.769, $p$ = 2.6x10$^{-6}$).

**Table S1 – Long-term stability of activation across contrasts**

| Sample | Contrast | Mean | Median | SD | Min | Max |
|---|---|---|---|---|---|---|
| PMCP | Anticipation: Win > Baseline | 0.327 | 0.331 | 0.093 | 0.049 | 0.835 |
| | Anticipation: Loss > Baseline | 0.234 | 0.229 | 0.107 | 0 | 0.824 |
| | Anticipation: Win > Loss | 0.0501 | 0.0244 | 0.0726 | 0 | 0.786 |
| | Feedback: Win > Baseline | 0.219 | 0.22 | 0.101 | 0 | 0.668 |
| | Feedback: Loss > Baseline | 0.246 | 0.244 | 0.101 | 0 | 0.811 |
| | Feedback: Win > Loss | 0.0768 | 0.0695 | 0.0684 | 0 | 0.285 |
| | Feedback: Win > Neutral | 0.102 | 0.103 | 0.0695 | 0 | 0.328 |
| | Feedback: Loss > Neutral | 0.011 | 0 | 0.0323 | 0 | 0.195 |
| PGS-E | Anticipation: Win > Baseline | 0.116 | 0.108 | 0.0705 | 0 | 0.578 |
| | Anticipation: Loss > Baseline | 0.124 | 0.122 | 0.069 | 0 | 0.475 |
| | Anticipation: Win > Loss | 0.051 | 0.0448 | 0.0446 | 0 | 0.208 |

Long term stability of reward activation across contrasts in the Pitt Mother Child Project (PMCP) study and the Pittsburgh Girls Study (PGS-E), using the Schaefer-400 cortical parcellation and the Harvard-Oxford subcortical parcellation. A stability of ‘0’ indicates that the stability is smaller than could be estimated, and thus defaulted to 0.

**Table S2 – Stability of activation across networks during reward anticipation**

| Sample | Network | Mean | Median | SD | Min | Max |
|---|---|---|---|---|---|---|
| PMCP | Limbic | 0.252 | 0.261 | 0.0679 | 0.109 | 0.429 |
| | Control | 0.299 | 0.31 | 0.0848 | 0.0867 | 0.445 |
| | Default | 0.298 | 0.301 | 0.0778 | 0.103 | 0.538 |
| | Attention | 0.355 | 0.34 | 0.0802 | 0.187 | 0.513 |
| | Reward | 0.255 | 0.262 | 0.105 | 0.049 | 0.413 |
| | Salience | 0.369 | 0.383 | 0.089 | 0.142 | 0.555 |
| | Somatomotor | 0.384 | 0.388 | 0.0928 | 0.181 | 0.835 |
| | Visual | 0.324 | 0.332 | 0.0727 | 0.0857 | 0.458 |
| PGS-E | Limbic | 0.05 | 0.0372 | 0.0568 | 0 | 0.202 |
| | Control | 0.15 | 0.151 | 0.0639 | 0.0236 | 0.333 |
| | Default | 0.138 | 0.132 | 0.0688 | 0.0129 | 0.578 |
| | Attention | 0.117 | 0.115 | 0.0597 | 0 | 0.261 |
| | Reward | 0.0614 | 0.0533 | 0.0381 | 0 | 0.132 |
| | Salience | 0.105 | 0.0916 | 0.0649 | 0 | 0.24 |
| | Somatomotor | 0.0989 | 0.0879 | 0.0727 | 0 | 0.27 |
| | Visual | 0.132 | 0.122 | 0.0669 | 0.011 | 0.373 |

Descriptors of the stability of reward activation across networks during the anticipation the Win>Baseline contrast, in the Pitt Mother Child Project (PMCP) study and the Pittsburgh Girls Study (PGS-E). Regions are defined using the Schaefer-400 cortical parcellation and the Harvard-Oxford subcortical parcellation. A stability of ‘0’ indicates that the stability is smaller than could be estimated, and thus defaulted to 0.

**Table S3 – Effect of network in the PMCP and PGS**

| Sample | Variable | B | SE | t | p |
|---|---|---|---|---|---|
| PMCP | (Intercept) | 0.252 | 0.016 | 15.808 | **$3.32X10^{-44}$** |
| | Control | 0.047 | 0.020 | 2.400 | **0.017** |
| | Default | 0.047 | 0.018 | 2.563 | **0.001** |
| | Attention | 0.103 | 0.020 | 5.182 | **$3.47X10^{-7}$** |
| | Reward | 0.004 | 0.024 | 0.159 | 0.874 |
| | Salience | 0.117 | 0.020 | 5.916 | **$7.02X10^{-9}$** |
| | Somatomotor | 0.129 | 0.018 | 7.031 | **$8.76X10^{-12}$** |
| | Visual | 0.072 | 0.019 | 3.793 | **$1.71X10^{-4}$** |
| PGS | (Intercept) | 0.050 | 0.012 | 4.123 | **$4.55X10^{-5}$** |
| | Control | 0.099 | 0.015 | 6.652 | **$9.43X10^{-11}$** |
| | Default | 0.085 | 0.014 | 6.127 | **$2.14X10^{-9}$** |
| | Attention | 0.067 | 0.015 | 4.418 | **$1.28X10^{-5}$** |
| | Reward | 0.011 | 0.018 | 0.638 | 0.524 |
| | Salience | 0.055 | 0.015 | 3.649 | **$2.98X10^{-4}$** |
| | Somatomotor | 0.049 | 0.014 | 3.484 | **$5.48X10^{-4}$** |
| | Visual | 0.082 | 0.015 | 5.623 | **$3.51X10^{-8}$** |

Results from linear regressions predicting region stability in the Pitt Mother Child Project (PMCP) study and the Pittsburgh Girls Study (PGS). Network-membership was dummy-coded and used as the predictive variables. The limbic network was the reference category, and thus is not shown in the table.

Figure S1 – Participant ages and time between study visits

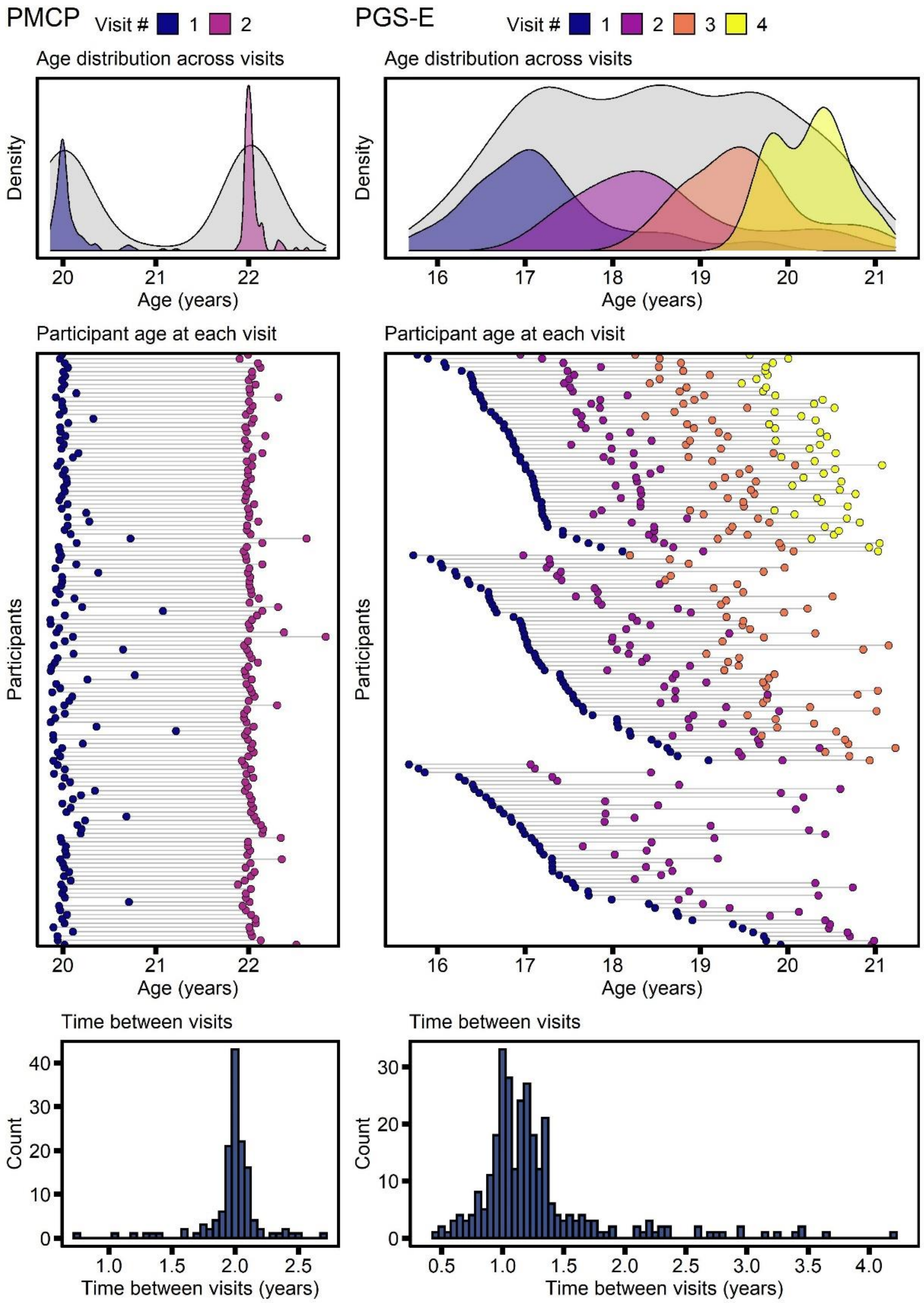


Distribution of participant age, across and within waves, in the Pitt Mother Child Project (PMCP) study and the Pittsburgh Girls Study (PGS).

Figure S2 – Monetary Task design

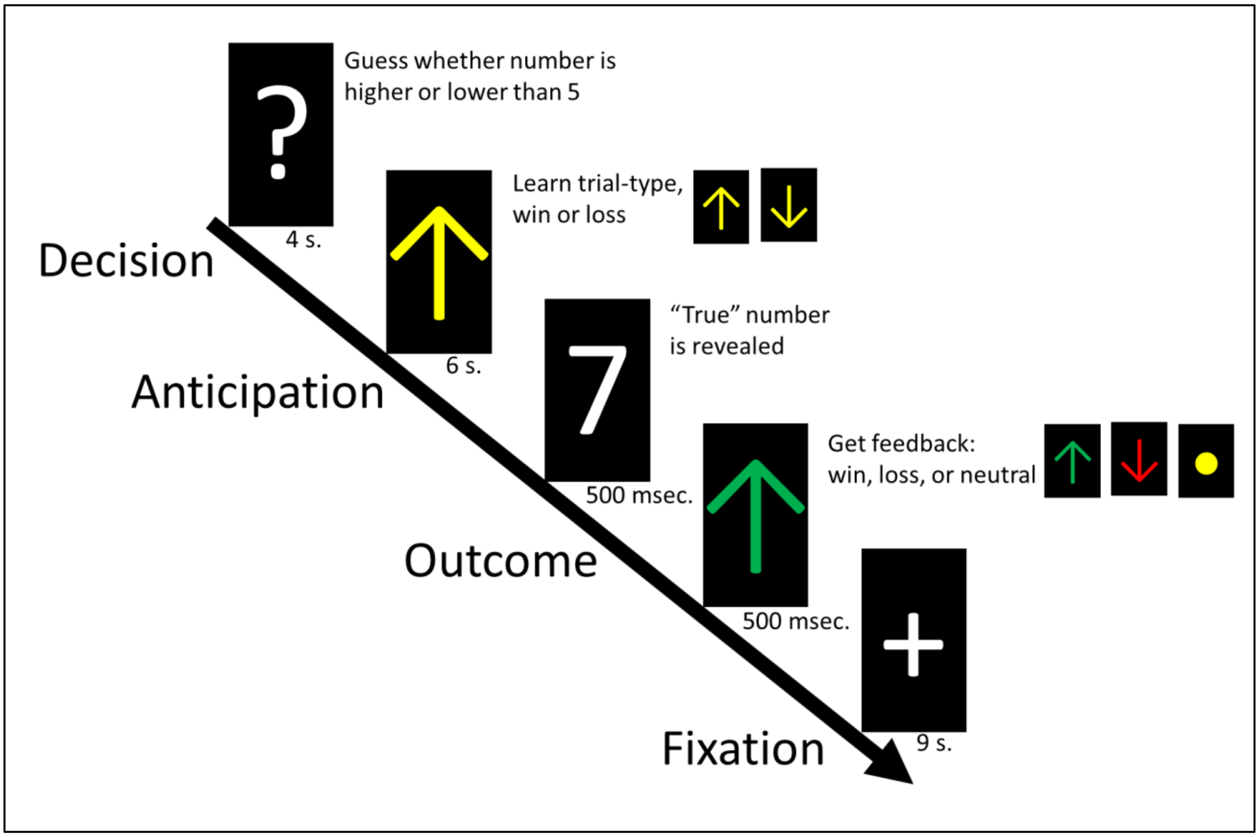


Schematic representation of the fMRI task design, see Methods. After Forbes et al. (Forbes et al., 2009)

Figure S3 – Reward Network from Neurosynth

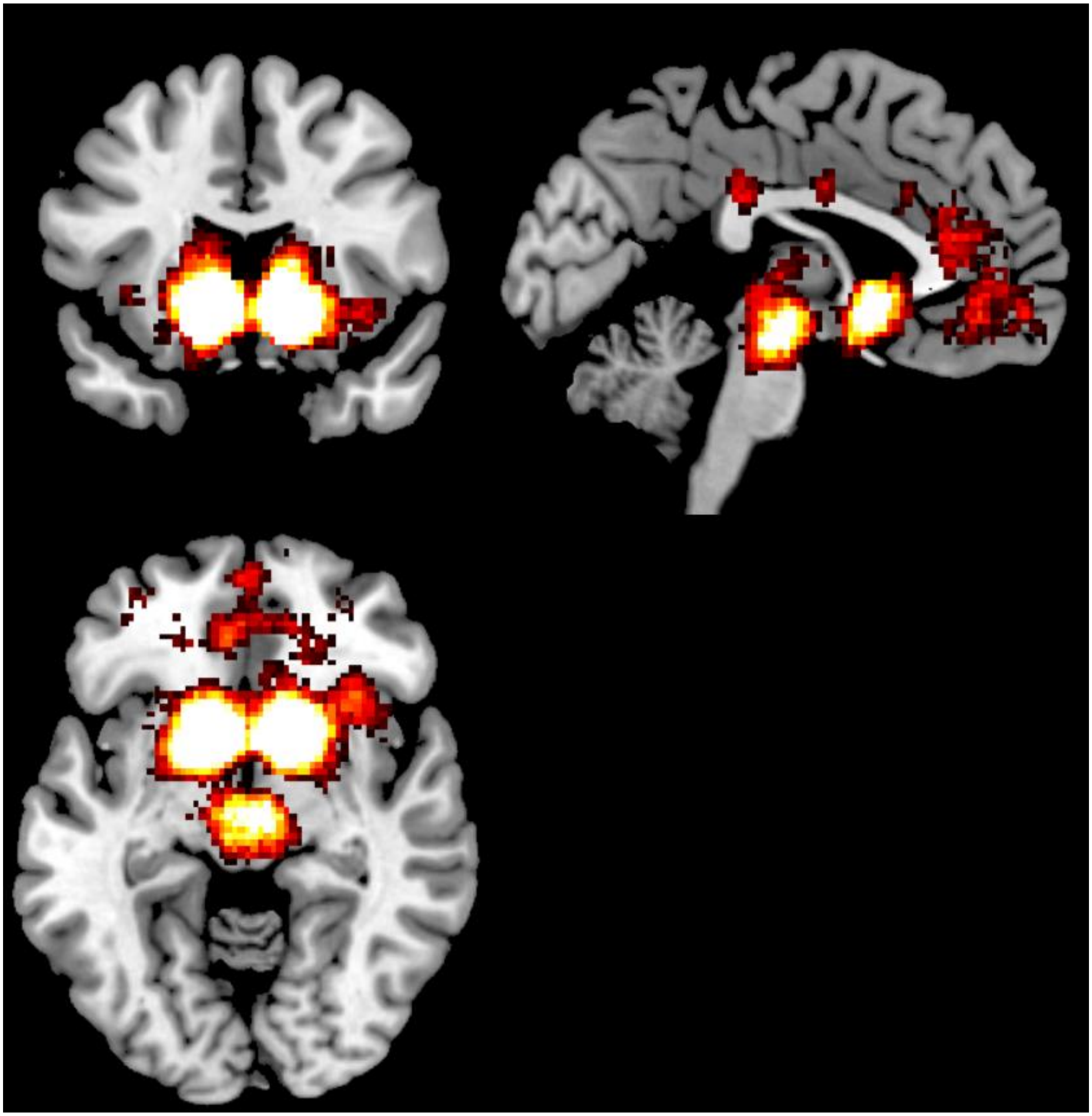

Results from Neurosynth (Yarkoni et al., 2011) used to select regions to include in the 'Reward' network. Results were accessed from https://neurosynth.org/analyses/topics/v5-topics-50/7. Across 14,371 studies, the 1,218 reward-tagged studies are more likely to report significant activation in these regions ($p<0.01$, FDR-corrected). Regions that likely to be reported in a 'reward' study include the caudate, putamen, nucleus accumbens, pallidum, amygdala, and thalamus, as well as portions of the orbitofrontal cortex, ventral medial prefrontal cortex, medial prefrontal cortex, insula, and the anterior cingulate.

Figure S4 – Stability does not differ across parcellations

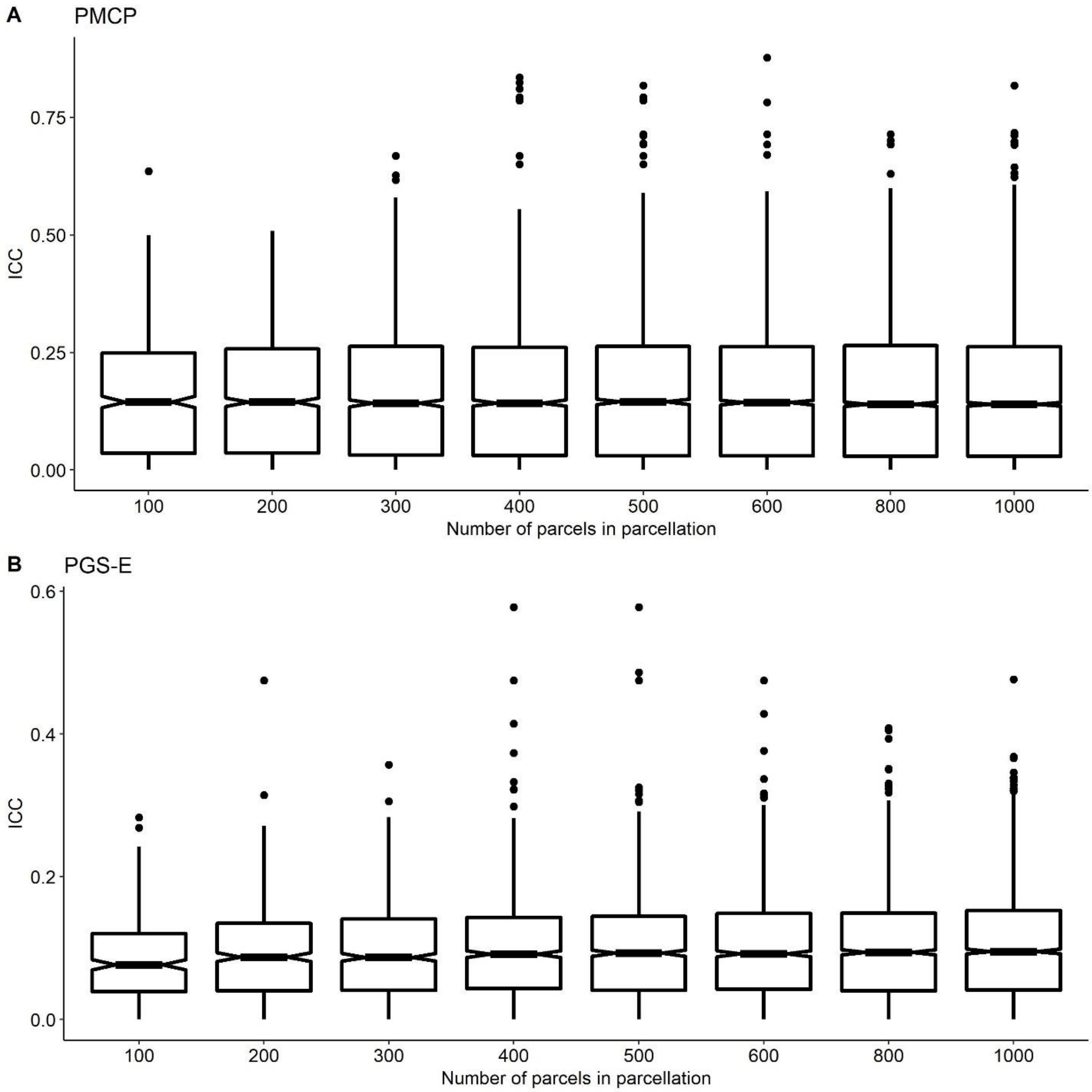


Stability of activation of cortical regions across the different cortical Schaefer parcellations (100-1,000 regions in each parcellation. (A) the Pitt Mother Child Project (PMCP) study and (B) the Pittsburgh Girls Study-Emotions Substudy (PGS-E). In the PGS-E the 100 and 200 parcellations are less stable than the others, there is no longer an effect of parcellation once those are removed (R=0.001, *p*=0.131).

Figure S5 – Contrast stability does not differ across parcellations

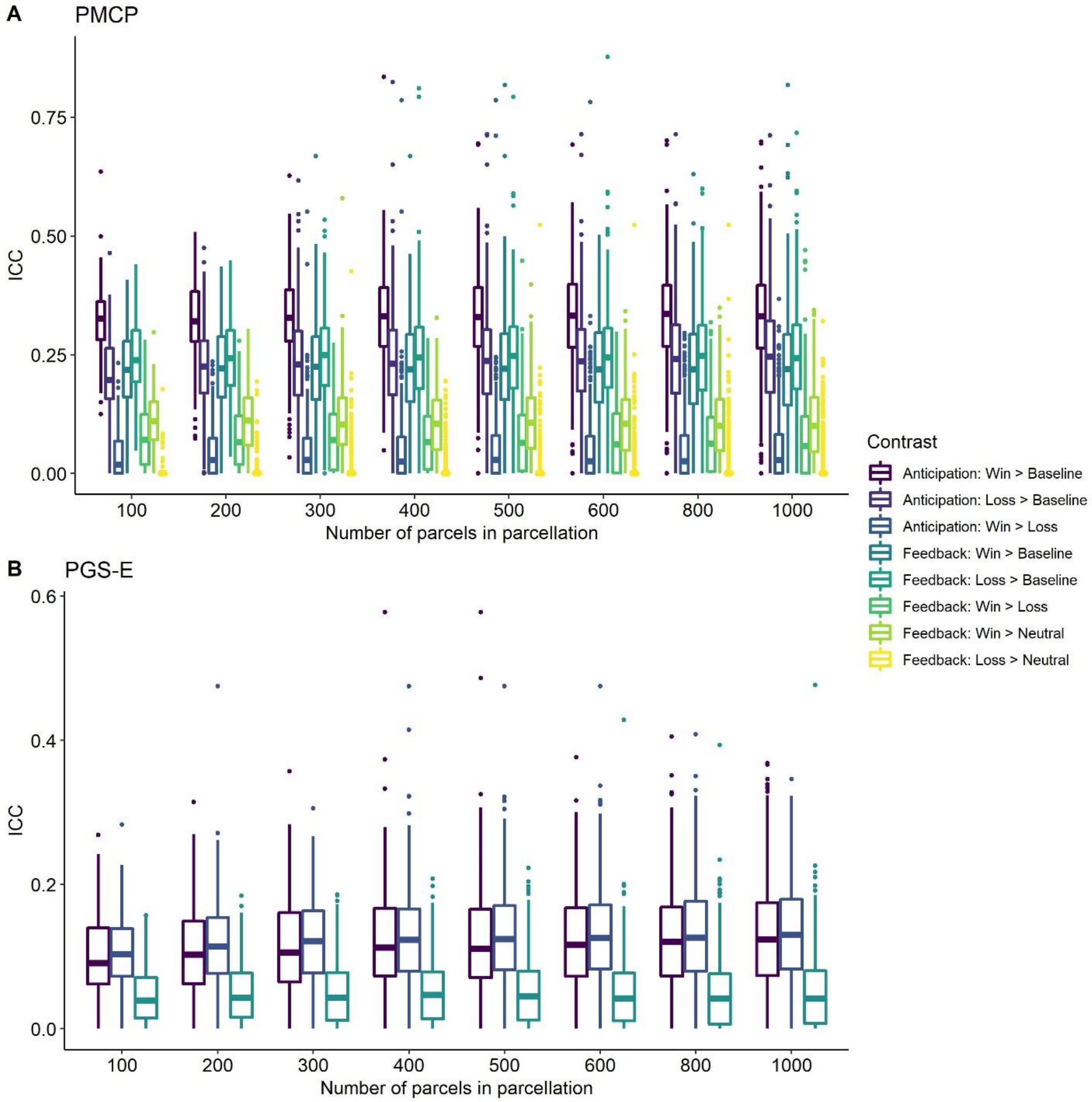


Stability of activation of cortical regions across all contrasts in all the cortical Schaefer parcellations (100-1,000 regions in each parcellation. (A) the Pitt Mother Child Project (PMCP) study and (B) the Pittsburgh Girls Study (PGS).
